# Milk Fat Globule Membrane-Containing Protein Powder Promotes Fitness in *Caenorhabditis elegans*

**DOI:** 10.1101/2024.06.24.600324

**Authors:** Miina Pitkänen, Olli Matilainen

**Affiliations:** The Molecular and Integrative Biosciences Research Programme, Faculty of Biological and Environmental Sciences, University of Helsinki, Helsinki, Finland

**Keywords:** milk fat globule membrane (MFGM), hydrolysated milk protein, *C. elegans*, fitness, innate immunity, cathepsin B

## Abstract

Milk-derived peptides and milk fat globule membrane (MFGM) have gained interest as health-promoting food ingredients. However, the mechanisms by which these nutraceuticals modulate the function of biological systems often remain unclear. We utilized *Caenorhabditis elegans* to elucidate how milk-derived Protein powders rich in MFGM, previously used in a clinical trial, affect the physiology of this model organism. Our results demonstrate that Protein powders do not affect lifespan but promote the fitness of the animals. Surprisingly, gene expression analysis revealed that Protein powders decrease the expression of genes functioning on innate immunity, which also translates into reduced survival on pathogenic bacteria. One of the innate immunity-associated genes showing reduced expression upon Protein powder supplementation is *cpr-3*, the homolog of human cathepsin B. Interestingly, knockdown of *cpr-3* enhances fitness, but not in Protein powder-treated animals, suggesting that protein powders contribute to fitness by downregulating the expression of this gene. In summary, this research highlights the value of *C. elegans* in testing the biological activity of food supplements and nutraceuticals. Furthermore, this study should encourage investigations into whether milk-derived peptides and MFGM mediate their beneficial effects through the modulation of cathepsin B expression in humans.

## 1. Introduction

In addition to exercise, nutrition plays a central role in maintaining health and fitness, which is crucial for both societal welfare and individual well-being. Therefore, food supplements and nutraceuticals have become increasingly popular for complementing the diet. These products have potential to fill nutritional gaps and thereby prevent chronic diseases by providing essential vitamins, minerals, and other bioactive compounds [1]. Milk-derived bioactive peptides, which are naturally present in milk or can be generated through the hydrolysis of native proteins, have garnered attention due to their diverse health benefits and potential applications in functional foods, dietary supplements, and pharmaceuticals. For example, these peptides have been shown to promote cardiovascular health, possess immunomodulatory properties, and improve metabolic health [2,3]. In addition to milk-derived peptides, milk fat globule membrane (MFGM) is another bioactive component found in milk. MFGM, which encloses milk fat to globules, has a complex trilayer structure mainly composed of polar lipids, cholesterol, various proteins and glycoproteins [4]. MFGM is recognized for its potential health benefits, especially in infants, as it promotes, for example, the maturation of the gut and the immune system, as well as boosts the cognitive development [5,6]. Studies conducted on older subjects have found that MFGM-enriched milk improves episodic memory [7], while MFGM supplementation combined with exercise may be beneficial in improving walking speed and other walking parameters, such as step length [8].

Recently, it was reported how Protein powder (with partially hydrolyzed protein) and daily snack rich in both milk-derived peptides and MFGM affect the physical performance of older women [9]. This study found that these dietary interventions improve the physical performance based on the Short Physical Performance Battery score [9]. Building on this study, we utilized the widely used model organism, *Caenorhabditis elegans*, to further investigate the effects of MFGM-containing protein powders on a multicellular organism. Due to its small size, ease of culture, and short life span, *C. elegans* is an excellent model to study how different interventions affect an organism’s physiology throughout its life cycle. Notably, despite being a 1 mm long nematode, at least 83% of *C. elegans* proteome has human homologous genes [10]. Moreover, as *C. elegans* conserves the key signaling pathways regulating health- and lifespan across eukaryotes, it is emerging model in food and nutrition research, as well as in drug discovery [11,12].

## 2. Materials and Methods

### 2.1 Protein powders

The production of Protein powders has been described earlier [9]. Shortly, lactose free, protein-hydrolyzed butter milk powder was produced from ultrafiltered lactose free buttermilk concentrate. Part of monosaccharides were removed by ultrafiltration, and proteins and residual fats were concentrated. After ultrafiltration, proteins were partially hydrolyzed by enzymatic hydrolysis. The lactose content of the final protein concentrate was <0.01 %. Protein hydrolysis was undertaken according to patent EP 2632277B1 [13] and as described previously [14]. For experiments in this study, Protein powders were mixed with water to achieve a final concentration of 1 mg/ml. Notably, Protein powders are not fully soluble in water. Therefore, to increase their solubility, they were ground with a plastic pestle in 500 μl of water in a 1.5 ml Eppendorf tube before diluting them to a concentration of 1 mg/ml. Protein powder solutions were spread on agar plates (200 µl on a 3 cm plate, 1 ml on a 6 cm plate, and 2 ml on a 10 cm plate). Water was used as a control in all experiments. After the plates dried, they were seeded with bacteria.

### 2.2 C. elegans maintenance

The *C. elegans* N2 (Bristol) strain was used in all experiments. *C. elegans* were maintained on NGM plates (peptone, P4963, Merck; agar, A4550, Merck; NaCl, 746398, Merck). Except PA14 assay and *cpr-3* RNAi experiments, Animals were kept on *E. coli* OP50 or *E. coli* HT115 carrying the empty vector (EV, control vector for RNAi). When using the HT115 (EV), bacterial culture was prepared according to the RNAi protocol described earlier (including the supplementation of IPTG) [15]. Unless otherwise mentioned, all experiments were performed at 20°C.

### 2.3 Lifespan analyses

Lifespan experiments were performed on *E. coli* OP50 or *E. coli* HT115 carrying an empty vector (EV). Lifespan experiments were initiated by allowing gravid hermaphrodites (P0 generation) to lay eggs on 6 cm NGM agar plates, and the F1 generation was scored for lifespan. Alternatively, animals were bleached and allowed to hatch overnight in M9 before plating L1 larvae onto 6 cm NGM agar plates. These two alternative methods to initiate lifespan did not affect the conclusions drawn from the experiments. At the L4 larval stage, animals were transferred to 3 cm NGM agar plates containing 5-Fluorouracil (5-FU) (10 µM) (Merck, #F6627) to prevent progeny production. Animals with an exploded vulva or that crawled off the plate were censored. Animals were counted as dead if there was no movement after being poked with a platinum wire. Lifespans were checked every 1-3 days. Animals were maintained on protein powder plates throughout the entire experiment. Mean lifespan ± standard error (SE) is reported in the Supplementary Materials file 1, Table S1.

### 2.4 Activity measurement

Animals were synchronized by bleaching and plated as L1 larvae on 10 cm NGM agar plates seeded with *E. coli* OP50 or *E. coli* HT115 (EV), which were kept at 20°C. At the L4 larval stage, animals were transferred to 10 cm NGM agar plates containing 5-FU (10 µM) (Merck, #F6627) to prevent progeny production. Activity was measured on days 2 and 4 of adulthood. Animals were transferred to new plates after day 2 of adulthood. For activity measurement, 10 animals were placed in a single well of a 96-well plate containing 100 µl of M9 solution. Activity was measured for two hours using wMicroTracker (InVivo Biosystems).

### 2.5 RNA sequencing

Animals were synchronized by bleaching and plated as L1 larvae on control or Protein powder 2-containing 10 cm NGM agar plates seeded with *E. coli* OP50. At the L4 larval stage, animals were transferred to 10 cm NGM agar plates containing 5-FU (10 µM) (Merck, #F6627) to prevent progeny production. Animals were collected at day 2 of adulthood (three biological replicates for both strains) and frozen in liquid nitrogen. Total RNA was extracted with TRIzol Reagent (ThermoFisher Scientific, #15596018). Samples were sent to Novogene for library construction, quality control, sequencing, and data analysis. In short, mRNA was purified from total RNA using poly-T oligo-attached magnetic beads. After the fragmentation, the first strand cDNA was synthesized using random hexamer primers, followed by the second strand cDNA synthesis. The library preparations were sequenced on an Illumina platform. Paired-end clean reads were mapped to the reference genome using HISAT2 software [16]. FeatureCounts [17] was used to count the read numbers mapped of each gene. Differential expression analysis between two conditions (three biological replicates per condition) was performed using DESeq2 [18]. Genes with p-value < 0.05 and log2FoldChange > 0 found by DESeq2 were assigned as differentially expressed. Differentially expressed genes can be found from Supplementary Materials file GO enrichment analysis was done using clusterProfiler [19]. The RNA-seq data are available in the Gene Expression Omnibus (GEO) database repository (GSE270138).

### 2.6 Quantitative RT-PCR (qRT-PCR)

Animals were synchronized by bleaching and plated as L1 larvae on 10 cm NGM agar plates. Plates were kept at 20°C. Animals were collected at the L4 larval stage or on day 2 of adulthood and frozen in liquid nitrogen. Animals that were collected on day 2 of adulthood were transferred to plates containing 5-FU (10 µM) (Merck, #F6627) at the L4 larval stage to prevent progeny production. TRIzol Reagent (ThermoFisher Scientific, #15596018) was used to extract RNA. cDNA synthesis was performed with the QuantiTect Reverse Transcription Kit (Qiagen, #205313), and qRT-PCR reactions were run with the HOT FIREPol SolisGreen qPCR Mix reagent (Solis BioDyne, #08-46-00001) using the CFX384 or CFX Opus 384 machine (Bio-Rad). qRT-PCR data were normalized to the expression of *cdc-42* and *pmp-3*. qRT-PCR oligos used in this study are provided in Supplementary Materials file 1, Table S2. qRT-PCR experiments were performed with three biological replicates, with three technical replicates for each biological replicate. All qRT-PCR experiments were performed at least twice.

### 2.7. Pseudomonas aeruginosa assay

*Pseudomonas aeruginosa* assays were performed as described earlier [20] with minor modifications. 200 μl of Protein powder solutions (1 mg/ml) were added to 3 cm NG plates with 10 µM 5-FU (Merck, #F6627). After the plates had dried, they were seeded with 3 µl of an overnight-grown *Pseudomonas aeruginosa* (PA14) suspension and incubated at 37°C for 24 hours. 20 μl of 2% SDS was added to the edges of the plate to prevent the escape of the animals. *C. elegans* were grown on 6 cm NGM agar plates supplemented with 1 ml of Protein powder solutions (1 mg/ml) and seeded with OP50. Animals were transferred to PA14 plates at the L4 stage and incubated at 25°C. Animals were scored daily for survival based on their ability to respond to touch. Animals that crawled off the plate were censored. Mean lifespan on PA14 ± standard error (SE) is reported in the Supplementary Materials file 1, Table S1.

### 2.8 RNA interference (RNAi)

*cpr-3* RNAi clone was taken from the Ahringer RNAi library. RNAi was performed using the feeding protocol described earlier [15]. Animals were kept on *cpr-3* RNAi during the whole experiment.

### 2.9. Statistical analysis

Statistical analyses for motility and qRT-PCR data were carried out in GraphPad Prism. qRT-PCR data represent the mean of three biological replicates ± standard deviation (SD). Statistical details can be found in the figures and figure legends. Statistical calculations for lifespan experiments were carried out in R using the Cox-proportional hazard regression analysis. Statistical details for the lifespan data can be found in Supplementary Materials file 1, Table S1.

## 3. Results

### 3.1 Protein powders improve C. elegans motility

In this study we used two protein powders (Protein powder 1 and 2), which were generated using the same recipe but are from two independent batches (see Methods for infromation on Protein powder production). These Protein powders differ slightly in their MFGM content, as it is 13,5 mg/g and 14 mg/g in Protein powder 1 and Protein powder 2, respectively. Regarding the concentration of Protein powders used in the experiments, it has been shown that single amino acids extend *C. elegans* lifespan in liquid S-medium when supplemented with concentrations of 1-10 mM [21]. 10 mM of serine, which shows the most prominent effect on lifespan [21], equals approximately the concentration of 1 mg/ml. Since a single amino acid has a robust effect on lifespan at this concentration, we decided to investigate whether Protein powders affect physiology with similar concentrations. For this, we mixed protein powders with water to achieve a final concentration of 1 mg/ml and spread the solutions on NGM agar plates (see Methods for details).

At first, we tested whether Protein powders affect lifespan. For lifespan experiments, we used the *E. coli* B strain OP50, which is probably the most often used bacterial strain in *C. elegans* maintenance and experiments, as a food source. In addition, we utilized the *E. coli* K-12 strain HT115, which is another commonly utilized bacterial strain in *C. elegans* research, particularly in RNAi experiments. In this experiment we employed HT115 bearing the empty vector (EV), which serves as a control in RNAi experiments. Notably, HT115 provides a healthier diet compared to OP50, as it not only improve the host’s response to oxidative, heat, or pathogenic stress [22] but also leads to a longer lifespan [23–25]. In the lifespan assays performed at 20°C, we found that Protein powders do not affect lifespan on either diet (Figure 1a-b, Supplementary Materials file 1, Table S1). In addition to the lifespan experiments conducted at 20°C, we also tested the effects of protein powders on the lifespan of animals kept at 25°C, a condition that induces mild heat stress. We conducted two independent experiments at this higher temperature. One experiment showed no statistical difference between the treatments, while the other experiment indicated that Protein powders induce a modest lifespan extension (Figure S1a, Supplementary Materials file 1, Table S1). Nevertheless, based on our data, it can be concluded that Protein powders do not affect lifespan.

**Figure 1.**
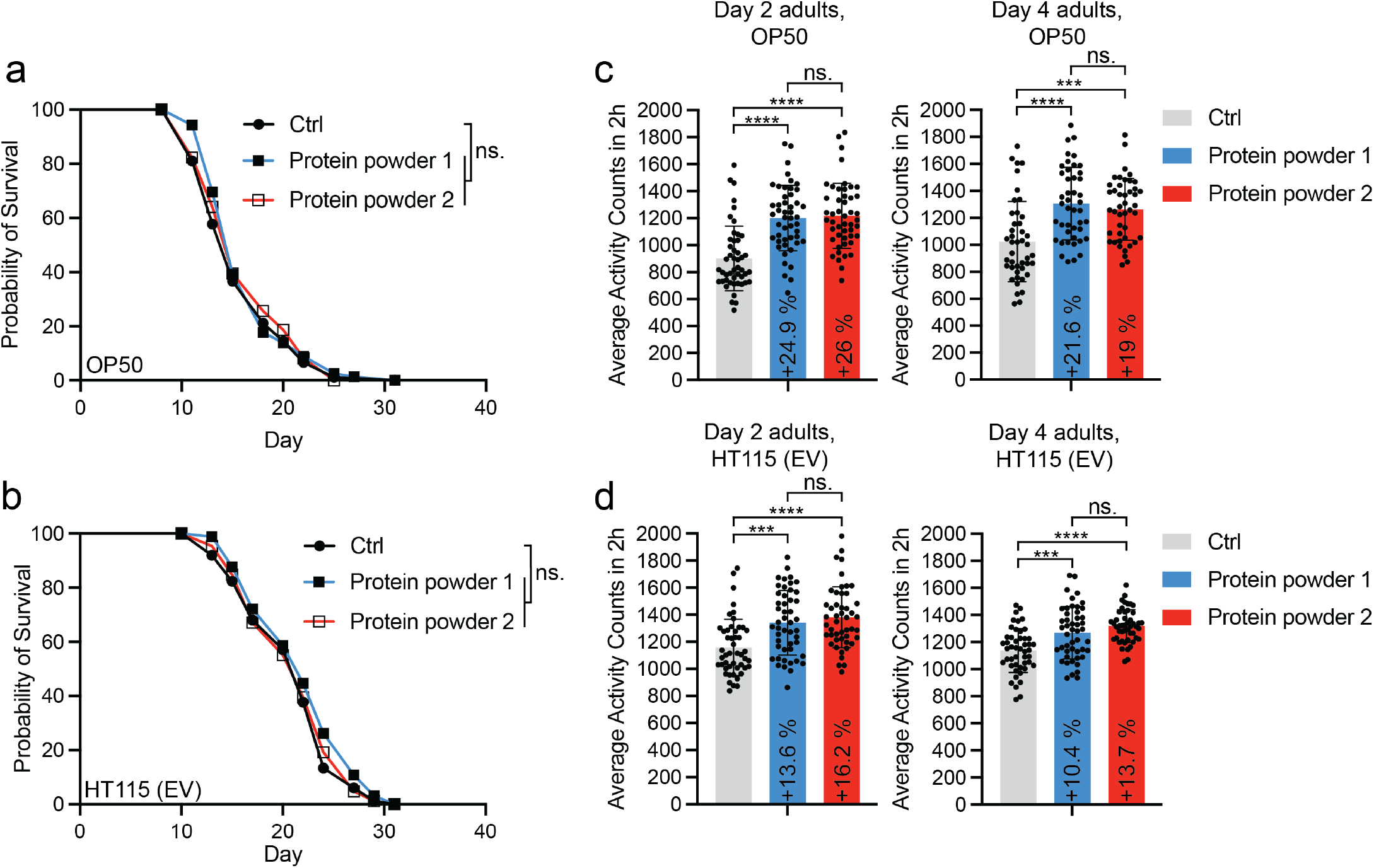
Protein powders do not affect *C. elegans* lifespan, but enhance motility. (**a**) Lifespan of OP50-fed and (**b**) HT115 (EV)-fed *C. elegans* on plates supplemented with Protein powder 1 or Protein powder 2. Lifespan statistics are reported in Supplementary Materials file 1, Table S1. (**c**) Motility of day 2 and day 4 adult (day 5 and day 7 from hatch, respectively), OP50-fed and (**d**) HT115 (EV)-fed *C. elegans* on plates supplemented with Protein powder 1 or Protein powder 2. Each dot represents a group of 10 animals (one well in a 96-well plate, n = 440 animals for day 4 adults on OP50, n = 480 animals for other conditions). Data are combined from two independent experiments (***p < 0.001, ****p < 0.0001, one-way ANOVA with Tukey’s test).

Importantly, lifespan does not always correlate with healthspan, and therefore many interventions that prolong lifespan also extend the period of frailty [26,27]. Therefore, we investigated whether Protein powders affect organismal health. For this, we used a thrashing assay in liquid, which measures *C. elegans* motility and is a widely used method to analyze the fitness of the animals [26,27]. In this assay, we placed animals in M9 solution and measured their swimming activity on a 96-well plate using WMictotracker (see Methods for details). Interestingly, when analyzing day 2 and day 4 adult animals (day 5 and day 7 from hatch, respectively) grown on OP50, we found that both Protein powder 1 and Protein powder 2 increase the motility of the animals (Figure 1c). Protein powders also increase the motility of animals grown on HT115 (Figure 1d), although their effect is not as pronounced as in experiments performed with OP50 (Figure 1c). Importantly, the HT115 diet alone leads to elevated motility compared to OP50-fed animals (Fig S1b), which may explain the finding that protein powders have a greater effect on animals maintained on OP50. Nevertheless, these data demonstrate that, although the protein powders do not affect lifespan, they promote the fitness of the animals.

### 3.2 Protein powders decrease the expression of genes related to innate immunity, and reduce the survival on pathogenic bacteria

To elucidate the mechanism by which protein powders promote fitness, we performed RNA-seq of OP50-fed day 2 adult *C. elegans* grown on Protein powder 2. Data analysis revealed that the expression of 1,210 genes is upregulated, and 1,077 genes are downregulated in Protein Powder 2-treated animals (Figure 2a, Supplementary Materials file 2). Interestingly, Gene Ontology (GO) enrichment analysis with differentially expressed genes revealed that downregulation of innate immunity-related processes show the most significant enrichment in Protein powder 2-treated animals (Figure 2b, Supplementary Materials file 2). Although the GO terms “*monocarboxylic acid metabolic process*” and “*carboxylic acid metabolic process*” also show strong enrichment among downregulated genes in Protein powder 2-treated animals (Figure 2b), the downregulation of innate immunity-related genes in three biological replicates is statistically more significant compared to the genes falling under the two abovementioned GO terms. For example, of the 22 most significantly downregulated genes, eight are under the innate immunity-related GO terms (Supplementary Materials file 2). In contrast, of the 106 most significantly downregulated genes, only three and four genes fall under the GO terms of “*monocarboxylic acid metabolic process*” and “*carboxylic acid metabolic process*,” respectively (Supplementary Materials file 2).

**Figure 2.**
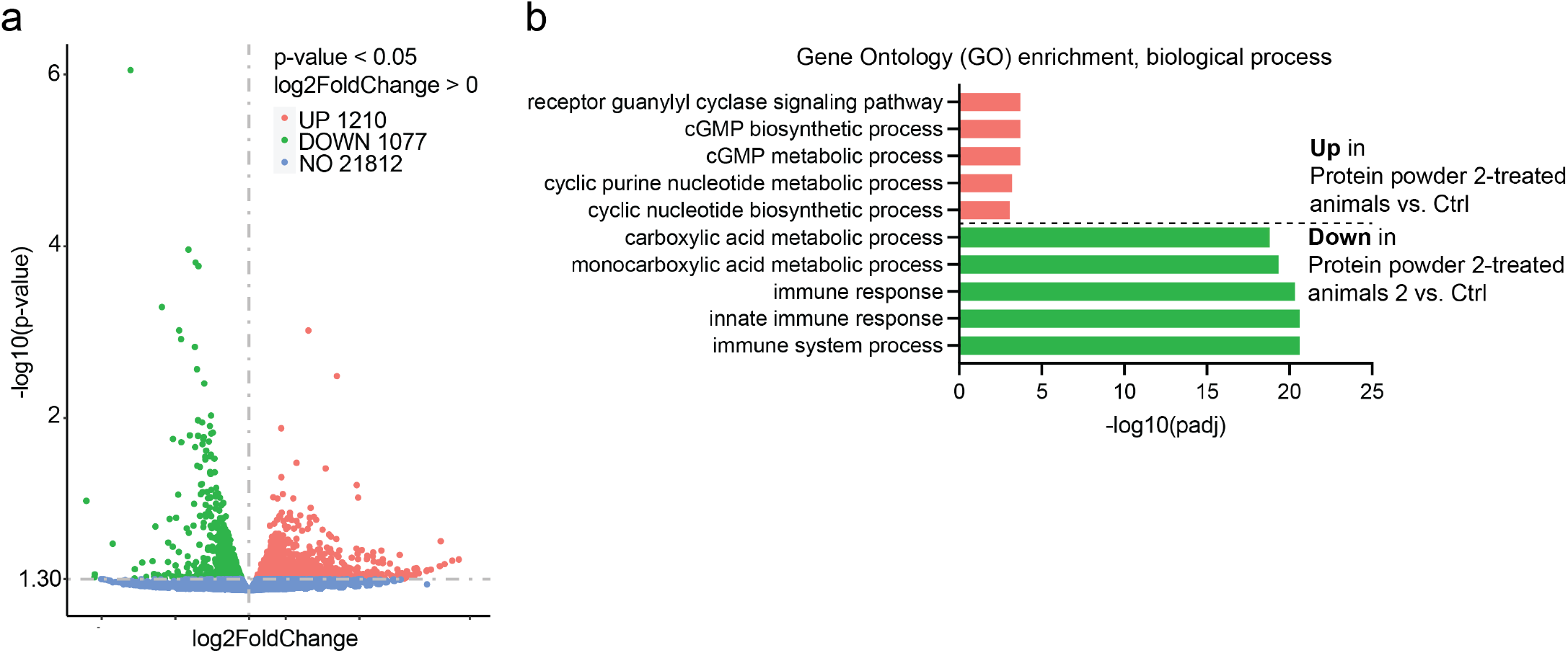
RNA sequencing of Protein powder 2-treated animals. (**a**) Volcano plot showing differentially expressed genes in of OP50-fed *C. elegans* treated with Protein powder 2 compared to control. (**b**) Enriched GO terms among up- and downregulated genes in Protein powder 2-treated animals. Lists of differentially expressed genes are shown in Supplementary Materials file 2.

Due to the downregulation of innate immunity-related genes in Protein powder 2-treated animals, we focused on this biological process. To validate the RNA-seq data, we performed qRT-PCR analysis on eight most significantly downregulated innate immunity-related genes from independent samples, including animals treated with Protein powder 1. These experiments revealed that both Protein powders reduce the expression of innate immunity genes in day 2 adult animals (Figure 3a). Based on these data, we hypothesized that Protein powder-treated animals may have an upregulated immune response earlier in development, which is then suppressed in post-developmental life stages. Therefore, we examined the expression of innate immunity genes at the L4 larval stage (day 3 from hatch, last larval stage before adulthood). In contrast to our expectations, most of the examined genes show reduced expression in L4 larvae as well (Figure S1c). In addition to OP50, we tested whether Protein powders affect the expression of innate immunity-related genes in animals fed with HT115 (EV), and found that protein powders also diminish the expression of innate immunity genes in day 2 adults on this diet (Figure 3b). Together, these data demonstrate that protein powders regulate the expression of innate immunity-related genes in a diet-independent manner. Moreover, as bacterial diet form *C. elegans* gut microbiome, our results demonstrate that Protein powders affect host physiology independently of these commensal bacteria.

**Figure 3.**
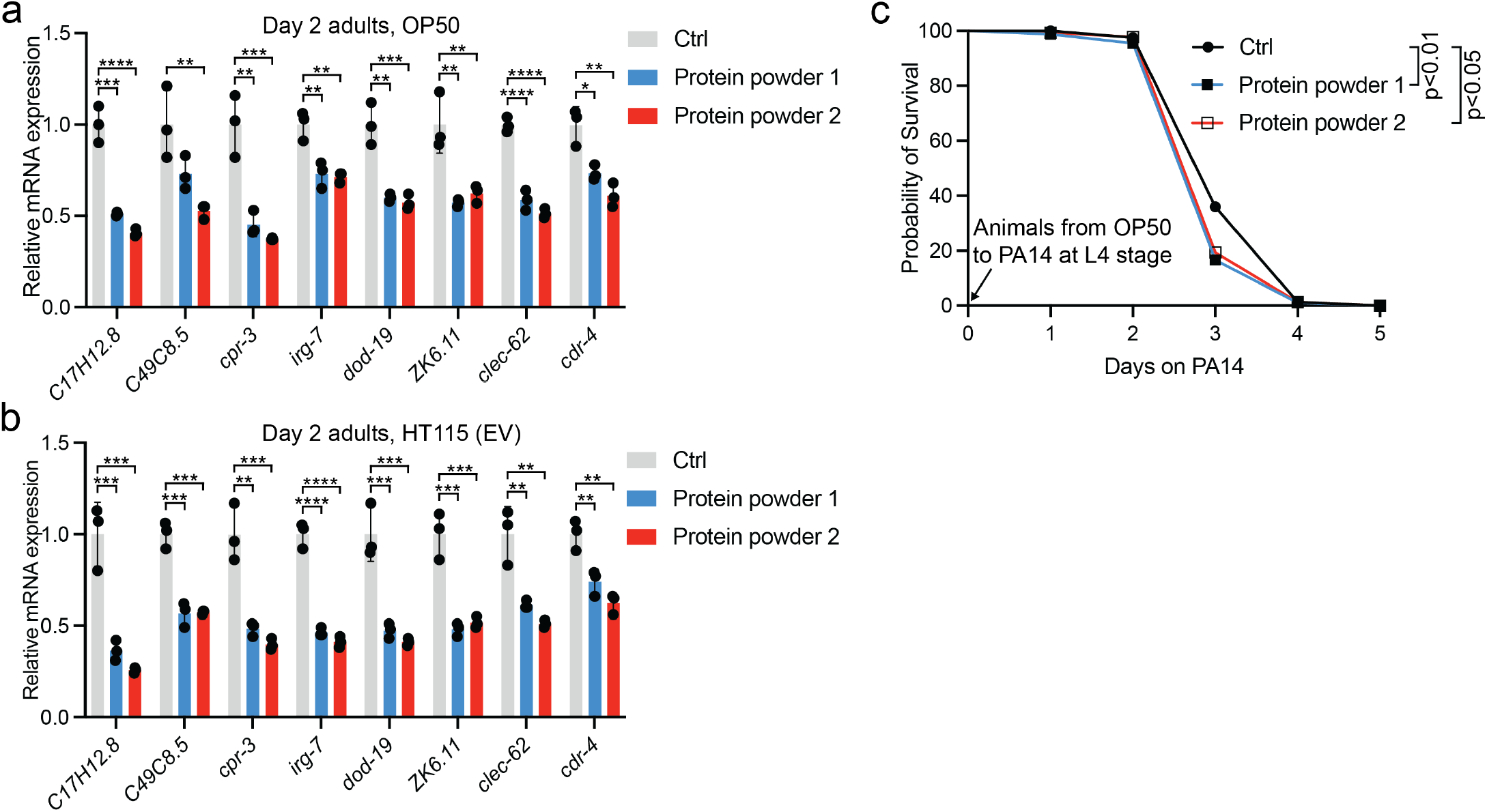
Protein powders reduce the expression of genes related to innate immunity. (**a**) qRT-PCR of selected innate immunity-related genes in OP50-fed and (**b**) HT115 (EV)-fed day 2 adult *C. elegans* grown on control or Protein powder-supplemented plates. Bars represent mRNA levels relative to control with error bars indicating mean ± SD of three biological replicates, each with three technical replicates (*p < 0.05, ** p < 0.01, ***p < 0.001, ****p < 0.0001, one-way ANOVA with Tukey’s test). (**c**) Survival of control- and Protein powder-treated animals on *Pseudomonas aeruginosa* (PA14). Statistics for PA14 assays are reported in Supplementary Materials file 1, Table S1.

As Protein powders reduce the expression of genes functioning in innate immunity (Figure 2b, Figures 3a-b), one obvious question arisises: do they impair survival on pathogenic bacteria? To investigate this, we performed a slow-killing assay with *Pseudomonas aeruginosa* strain PA14 [20]. Of the three independent experiments, one did not show differences in survival between control and Protein powder-treated animals (Supplementary Materials file 1, Table S1). However, in the two other experiment, Protein powders reduced survival on PA14 (Figure 3c, Supplementary Materials file 1, Table S1). These data suggest that, reflecting the decreased expression of genes associated with innate immunity, Protein powders may slightly suppress the innate immune response against pathogenic bacteria.

### 3.3 Downregulation of cpr-3 promotes fitness

Since Protein powders decrease the survival on pathogenic bacteria (Figure 3c, Supplementary Materials file 1, Table S1), we hypothesized that it is a trade-off for enhanced fitness (Figures 1c-d). In other words, we asked whether downregulation of innate-immunity-related genes promote fitness. Notably, one of the most significantly downregulated gene in Protein powder-treated animals is *cpr-3* (Figures 3a-b, Figure S1c,Supplementary Materials file 2). *cpr-3* is a homolog for cathepsin B (CTSB), a lysosomal cysteine protease [28]. Importantly, cathepsin B has been associated with many pathologies [29]. For example, clinical findings from multiple studies have shown that CTSB levels are increased in many neurologic conditions, including several neurodegenerative diseases such as Alzheimer’s disease and traumatic brain injury [30]. Similarly, as in humans, CTSB levels are increased in animals modeling neurologic disorders, and as shown in 12 studies, its deletion leads to significant improvements in behavioral deficits and neuropathology in these animal models [30]. Furthermore, CTSB has a role in the development of cancer as well as in conditions such as lung and cardiovascular disorders [31]. Based on these reports, we examined how *cpr-3* RNAi affects *C. elegans* fitness. Strikingly, we found that *cpr-3* knockdown leads to a significant increase in the motility of both day 2 and day 4 adult *C. elegans* (Figure 4a). Notably, *cpr-3* RNAi does not further increase the motility of Protein powder 2-treated animals (Figure 4a), indicating that these interventions promote fitness through the same mechanism.

**Figure 4.**
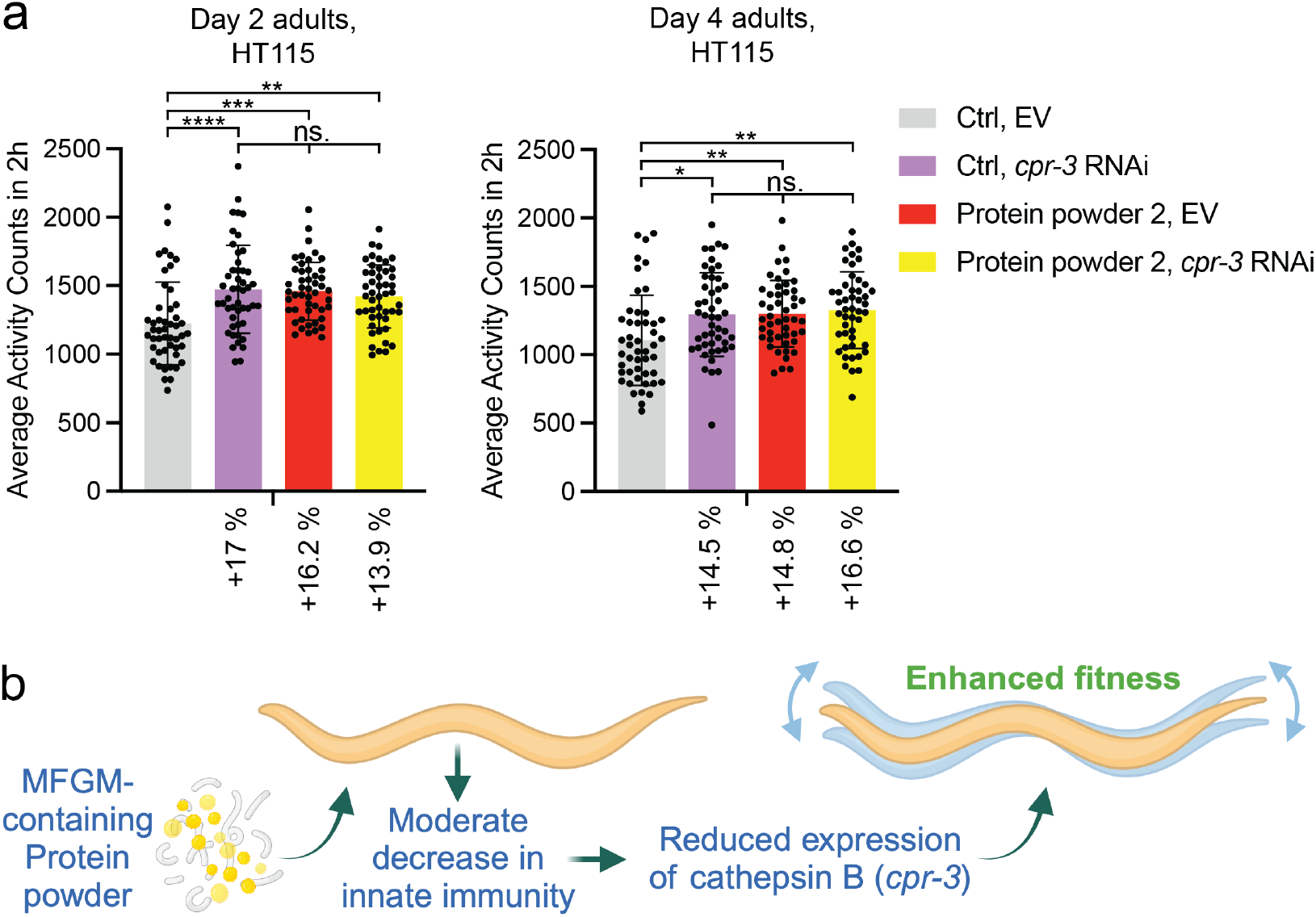
Knockdown of Protein powder-regulated cathepsin B enhances motility. (**a**) Motility of day 2 and day 4 adult, control (EV) or *cpr-3* RNAi-treated animals grown on control or Protein Powder 2-supplemented plates. Each dot represents a group of 10 animals (one well in a 96-well plate, n = 480 animals for all conditions). Data are combined from two independent experiments (*p < 0.05, **p < 0.01, ***p < 0.001, ****p < 0.0001, one-way ANOVA with Tukey’s test). (**b**) Model based on data presented in this study. Illustration was created with BioRender.com.

## 4. Discussion

We show here that MFGM-containing Protein powders promote fitness, which is, at least partly, due to the downregulation of *cpr-3* expression (Figure 4b). Interestingly, Protein powders uncouple fitness from lifespan, as they do not affect longevity (Figures 1a-b). These data indicate that their effect on fitness (Figures 1c-d) is not mediated through the known lifespan-modulating mechanisms. Furthermore, the finding that Protein powders enhance fitness on both *E. coli* OP50 and HT115 is important, as it indicates that the effect is independent of both diet and gut microbiome. To highlight the influence of diet/microbiome in *C. elegans* health- and lifespan experiments, metformin, a widely used drug for type 2 diabetes, has been shown to promote longevity on OP50, but not on the healthier HT115 strain [32]. In this respect, considering that MFGM may be beneficial in the treatment of obesity and the associated type 2 diabetes [33,34], MFGM-containing Protein powders could be additive to the effects of metformin in the aforementioned conditions.

With regard to the finding that *cpr-3* RNAi increases motility (Figure 4a), one might think that the phenotype is due to downregulation of CPR-3 function in muscle cells. However, data from a fluorescent reporter strain has shown that *cpr-3* is expressed in the pharynx, pharyngeal-intestinal valve, intestine, and rectal gland cells, with the intestine showing the strongest signal [35]. Therefore, in our experiments, it is likely that the Protein powder-induced reduction of *cpr-3* expression in the intestine promotes organismal fitness. This leads to the question, could a similar mechanism apply to humans? In humans, CTSB is not only ubiquitously expressed [36] (Human Protein Atlas, proteinatlas.org), but also secreted. Interestingly, running-induced systemic CTSB secretion from muscles induce beneficial cognitive effects [37], whereas CTSB secretion from many other cell types can have aggravating effects (for example through the modulation of extracellular matrix (ECM) [38]), especially under pathological conditions [31,39,40]. Thus, together with the physiological condition, the source tissue may affect whether the secreted CTSB is beneficial or not. Notably, adipose tissue has been shown to be one of the tissues that secrete CTSB [41]. Given that the intestine, which expresses *cpr-3* at high level [35], is the primary adipose tissue in *C. elegans* [42], one intriguing hypothesis is that intestinal cells secrete CRP-3, which then modulate the ECM in, for example, muscle cells. Adipose tissue-secreted CTSB could have the same function in humans. In this respect, adipose tissue has been associated with the regulation of muscle function, as aging-associated adipose inflammation in obese people can lead to the fatty infiltration in skeletal muscles, which results in decreased muscle strength and functionality [43]. Nevertheless, further research is required to elucidate whether Protein powders modulate CTSB levels in humans.

In addition to *cpr-3*, the protein powders were also found to reduce the expression of many other genes related to innate immunity (Figure 2, Figures 3a-b, Figure S1c, Supplementary Materials file 2). Consistently with the gene expression data, Protein powders moderately reduce the survival on pathogenic PA14 (Figure 3c). We hypothesized above that this decrease in innate immunity is a trade-off for enhanced fitness (Figures 1c-d). Supporting this hypothesis, it has been shown that the allocation of energy toward immune function restricts physical growth in children and preadolescents [44,45]. Additionally, a meta-analysis of data from poultry suggest that organism has to make a trade-off between immune function and other fitness-enhancing traits [46]. Moreover, immune response restricts plant development, whereas aberrantly activated innate immunity is toxic to *C. elegans* [47], thus further underlining the conservation of trade-off between immunity and fitness.

Our observed effects of MFGM-containing Protein powders on *C. elegans* immunity are somewhat contradictory to the findings from humans, as numerous studies have reported that MFGM enhances immunity in infants [6,48], whereas milk-derived peptides have also shown promise as immunity-enhancing molecules [49]. In addition to the trade-off hypothesis presented above, there are two other possible explanations. Firstly, *C. elegans* and humans diverged during evolution, and unlike humans, *C. elegans* has not developed adaptive immunity. Therefore, this model organism cannot be used to study the effects of Protein powders on this specific branch of the immune system. Secondly, since we assessed the survival of *C. elegans* on PA14 at the adult stage, and considering that MFGM modulates the immunity of newborns [6,48], one explanatory factor for the opposing results could be the developmental stage when exposed to pathogens. On the other hand, milk-derived peptides and MFGM have also been shown to promote health by suppressing the production of multiple inflammation-associated proteins such as IL-1β, IL-6, and TNF-α (suppressed by MFGM [50,51]), as well as MCP-1 (suppressed by milk-derived peptides [52]), which raises the possibility that Protein powder-mediated suppression of immune response is beneficial in humans. Interestingly, inhibition of CTSB leads to reduced expression of all four aforementioned inflammatory genes [53]. Hence, considering that Protein powders reduce *cpr-3*/*CTSB* expression in *C. elegans* (Figures 3a-b, Figure S1c), it is possible that milk-derived peptides and MFGM modulate immunity through CTSB in mammals.

Together, the data presented here, which show that Protein powders promote *C. elegans* fitness, support the findings from a clinical study performed earlier [9]. Although these two studies were performed in organisms that separated in evolution hundreds of millions of years ago, similar results from both systems provide strong evidence that the tested MFGM-containing Protein powders possess beneficial biological activity. Finally, referring to the effect of Protein powders on cathepsin B (*cpr-3*) expression (Figures 3a-b, Figure S1c), the efficacy of almost twenty CTSB inhibitors has been tested in the treatment of various diseases, ranging from cancer to nervous system-associated maladies [29]. Thus, if Protein powders were found to decrease *CTSB* mRNA levels in humans, it would provide a complementary mechanism to target CTSB activity in disease.

## Supplementary Materials

Supplementary Materials file 1, Supplementary Materials file 2.

## Author Contributions

Conceptualization, O.M.; methodology, M.P. and O.M.; validation, M.P. and O.M.; formal analysis, M.P. and O.M.; investigation, M.P and O.M.; resources O.M.; data curation, M.P. and O.M.; writing—original draft preparation, M.P. and O.M.; writing—review and editing, M.P. and O.M.; visualization, O.M.; supervision, O.M.; project administration, O.M.; funding acquisition, O.M. All authors have read and agreed to the published version of the manuscript.

## Funding

This work was funded by Research Council of Finland.

## Data Availability Statement

The RNA-seq data generated during the current study are available in the Gene Expression Omnibus (GEO) database repository (GSE270138).

## Acknowledgments

The authors thank Valio Ltd. for providing Protein powders and Dr Susana Garcia (University of Helsinki) for sharing reagents. Some strains were provided by the CGC, which is funded by NIH Office of Research Infrastructure Programs (P40 OD010440). This work was supported by Research Council of Finland and University of Helsinki.

## Conflicts of Interest

The authors declare no conflicts of interest.

## Notes

https://www.ncbi.nlm.nih.gov/geo/query/acc.cgi?acc=GSE270138

### Competing Interest Statement

The authors have declared no competing interest.

