## Supplementary Materials file 1 for "Milk Fat Globule Membrane-Containing Protein Powder Promotes Fitness in *Caenorhabditis elegans*"

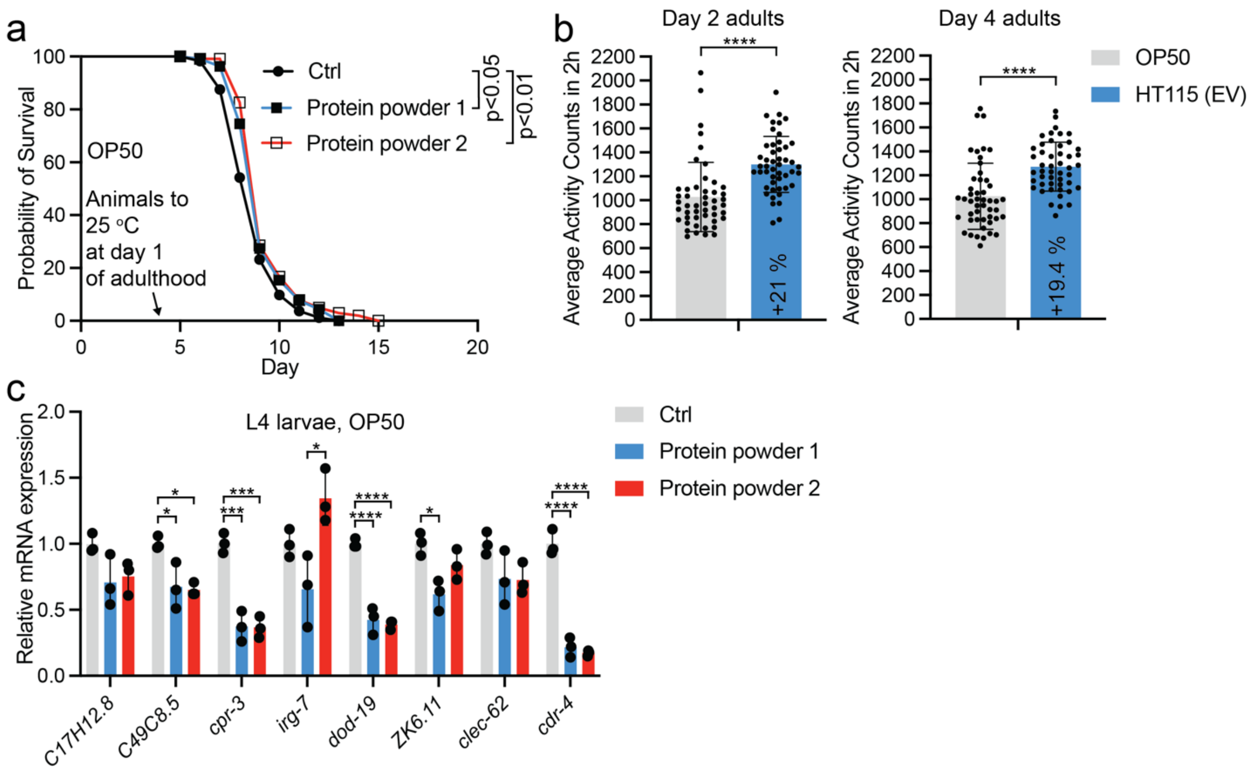


**Figure S1.** (**a**) Survival of OP50-fed *C. elegans* on control, Protein powder 1- and Protein powder 2-supplemented plates. Animals were transferred from 20°C to 25°C at day 1 of adulthood (4th day from hatch). Lifespan statistics are reported in Supplementary Information file 1, Table S1. (**b**) Motility of OP50- and HT115 (EV)-fed *C. elegans* on day 2 and day 4 of adulthood (day 5 and day 7 from hatch, respectively). Each dot represents a group of 10 animals (one well in a 96-well plate, n = 480 animals for all conditions). (**c**) qRT-PCR of selected innate immunity-related genes in OP50-fed, L4 larval stage (day 3 from hatch) *C. elegans elegans* grown on control or Protein powder-supplemented plates. Bars represent mRNA levels relative to control with error bars indicating mean ± SD of three biological replicates, each with three technical replicates (*p < 0.05, ***p < 0.001, ****p < 0.0001, one-way ANOVA with Tukey's test).

### Table S1. Individual replicates of *C. elegans* lifespan experiments

| Genotype and treatment | mean lifespan ± SE (days) | variation compared to control (%) | *P*-values against control | N |
|---|---|---|---|---|
| **Figure 1a** | | | | |
| N2 (OP50) Ctrl | 15.9 ± 0.45 | | | 91 |
| N2 (OP50), Protein powder 1 | 16.5 ± 0.45 | + 3.6 | 0.414 | 81 |
| N2 (OP50), Protein powder 2 | 16.2 ± 0.45 | + 1.9 | 0.621 | 88 |
| **Figure 1a** | | | | |
| N2 (OP50) Ctrl | 15.3 ± 0.44 | | | 85 |
| N2 (OP50), Protein powder 1 | 16.8 ± 0.41 | + 8.9 | 0.0807 | 67 |
| **Figure 1a** | | | | |
| N2 (OP50) Ctrl | 16.5 ± 0.47 | | | 90 |
| N2 (OP50), Protein powder 2 | 17.2 ± 0.54 | + 4.1 | 0.357 | 73 |
| **Figure 1a** | | | | |
| N2 (OP50) Ctrl | 14.8 ± 0.39 | | | |
| N2 (OP50), Protein powder 2 | 15.8 ± 0.57 | + 6.3 | | |
| **Figure 1b** | | | | |
| N2 (HT115, EV) Ctrl | 20.8 ± 0.48 | | | 85 |
| N2 (HT115, EV) Protein powder 1 | 20.8 ± 0.44 | + 0 | 0.673 | 85 |
| N2 (HT115, EV) Protein powder 2 | 21.5 ± 0.53 | + 3.3 | 0.492 | 68 |

| | | | | |
|---|---|---|---|---|
| **Figure 1b** | | | | |
| N2 (HT115, EV) Ctrl | 21.0 ± 0.49 | | | 83 |
| N2 (HT115, EV) Protein powder 1 | 22.0 ± 0.56 | + 4.5 | 0.135 | 67 |
| N2 (HT115, EV) Protein powder 2 | 21.2 ± 0.49 | + 0.9 | 0.765 | 84 |
| **Figure 1b** | | | | |
| N2 (HT115, EV) Ctrl | 20.3 ± 0.55 | | | 85 |
| N2 (HT115, EV), Protein powder 2 | 21.4 ± 0.55 | + 5.1 | 0.614 | 68 |
| **Figure S1a** | | | | |
| N2 (OP50) Ctrl, 25 ºC from day1 of adulthood | 8.78 ± 0.13 | | | 92 |
| N2 (OP50) Protein powder 1, 25 ºC from day1 of adulthood | 9.25 ± 0.14 | + 5.1 | 0.040294 | 93 |
| N2 (OP50) Protein powder 2, 25 ºC from day1 of adulthood | 9.44 ± 0.14 | + 7 | 0.003277 | 102 |
| **Figure S1a** | | | | |
| N2 (OP50) Ctrl, 25 ºC from day1 of adulthood | 9.66 ± 0.14 | | | 94 |
| N2 (OP50) Protein powder 1, 25 ºC from day1 of adulthood | 9.96 ± 0.17 | + 3 | 0.1824 | 98 |
| N2 (OP50) Protein powder 2, 25 ºC from day1 of adulthood | 10.08 ± 0.16 | + 4.2 | 0.0688 | 106 |
| **Figure 3c (days on PA14)** | | | | |
| N2 (PA14) Ctrl | 3.02 ± 0.029 | | | 155 |
| N2 (PA14) Protein powder 1 | 3.03 ± 0.032 | + 0.3 | 0.718 | 156 |
| N2 (PA14) Protein powder 2 | 2.95 ± 0.038 | - 2.3 | 0.725 | 155 |

| Figure 3c (days on PA14) | | | | |
|---|---|---|---|---|
| N2 (PA14) Ctrl | 3.35 ± 0.051 | | | 103 |
| N2 (PA14) Protein powder 1 | 3.12 ± 0.046 | - 6.9 | 0.004041 | 117 |
| N2 (PA14) Protein powder 2 | 3.1 ± 0.028 | - 7.5 | 0.021594 | 115 |
| **Figure 3c (days on PA14)** | | | | |
| N2 (PA14) Ctrl | 3.50 ± 0.068 | | | 134 |
| N2 (PA14) Protein powder 1 | 3.23 ± 0.046 | - 7.7 | 0.000319 | 143 |
| N2 (PA14) Protein powder 2 | 3.17 ± 0.042 | - 9.4 | 8.84e-06 | 146 |

### Table S2. Oligonucleotide sequences used in qRT-PCR

| Gene | Forward (5’ → 3’) | Reverse (5’ → 3’) |
|---|---|---|
| *cdc-42* | CTGCTGGACAGGAAGATTACG | CTCGGACATTCTCGAATGAAG |
| *pmp-3* | GTTCCCGTGTTCATCACTCAT | ACACCGTCGAGAAGCTGTAGA |
| *C17H12.8* | CACTGTCGATTGCTCACTCC | TCCGGTGCTGATGTATCCTT |
| *C49C8.5* | CGACTCAACTCTGCTTCTTGG | AGTGAAAAGAGCCATTCGAACT |
| *cpr-3* | GTGATTTCCGACCGAGTGTG | CTCCACCTGTTACTGCTCCA |
| *irg-7* | CGTGCCGGAACTATGTTCTG | ATCTCCGCTAGCTGTTCCTC |
| *dod-19* | AGTACCTCAGCCGACACTTC | AGCATCATCGAAATTGTAACGGA |
| *ZK6.11* | TGGCATATCTGTACGCTGGT | TGCTACCAAGGTCAACGAGA |
| *clec-62* | GCAAGAACAACACTCGCAAA | TGCTAACACCAGACGCCTTA |
| *cdr-4* | CGCTTCTGACTCGCTTTACA | TGCTCCAACACATCGGTAGT |
